# Bacterial Phage Tail-like Structure Kills Eukaryotic Cells by Injecting a Nuclease Effector

**DOI:** 10.1101/543298

**Authors:** Iara Rocchi, Charles Ericson, Kyle E. Malter, Sahar Zargar, Fabian Eisenstein, Martin Pilhofer, Sinem Beyhan, Nicholas J. Shikuma

## Abstract

Many bacteria interact with target organisms using syringe-like structures called Contractile Injection Systems (CIS). CIS structurally resemble headless bacteriophages and share evolutionarily related proteins such as the tail tube, sheath, and baseplate complex. Recent evidence shows that CIS are specialized to puncture membranes and often deliver effectors to target cells. In many cases, CIS mediate trans-kingdom interactions between bacteria and eukaryotes, however the effectors delivered to target cells and their mode of action are often unknown. In this work, we establish an *in vitro* model to study a CIS called <u>M</u>etamorphosis <u>A</u>ssociated <u>C</u>ontractile structures (MACs) that target eukaryotic cells. We show that MACs kill two eukaryotic cell lines, Fall Armyworm Sf9 cells and J774A.1 murine macrophage cells through the action of a newly identified MAC effector, termed Pne1. To our knowledge, Pne1 is the first CIS effector exhibiting nuclease activity against eukaryotic cells. Our results define a new mechanism of CIS-mediated bacteria-eukaryote interaction and are a first step toward understanding structures with the potential to be developed as novel delivery systems for eukaryotic hosts.

## INTRODUCTION

Bacteria interact with eukaryotic organisms with outcomes ranging from pathogenic to beneficial. One mechanism used by bacteria to interact with eukaryotes is through <u>C</u>ontractile Injection <u>S</u>ystems (CIS) (Taylor et al., 2018). CIS are evolutionarily related to the tails of bacteriophages (bacterial viruses) and are composed of an inner tube surrounded by a contractile sheath, capped with a tail spike and a baseplate complex. CIS can be classified into two types; Type 6 Secretion Systems (T6SS) and extracellular CIS (eCIS), also known as phage tail-like bacteriocins or tailocins. While T6SS reside within the bacterial cytoplasm and are anchored to the inner membrane of Gram-negative bacteria (Ho et al., 2014)(Böck et al., 2017), eCIS are released extracellularly by bacterial cell lysis and bind their target cell surface (Hurst et al., 2004; Shikuma et al., 2014; Yang et al., 2006). It has been speculated that eCIS may be an evolutionary intermediate between bacteriophage and T6SS (Büttner et al., 2016).

In both eCIS and T6SS, contraction of the sheath drives the inner tube and tail spike through the target cell membrane and both often deliver effectors to host cells. For example, an eCIS called *Photorhabdus* Virulence Cassettes (PVC) injects an effector causing actin condensation in insect hemocytes (Yang et al., 2006). In T6SS, a number of effectors are described that specifically target eukaryotic cells (Lien and Lai, 2017). The modes of action of these T6SS effectors include actin cross-linking in macrophages (Pukatzki et al., 2007), interaction with microtubules for invasion of epithelial cells (Sana et al., 2015), and disruption of the actin cytoskeleton of HeLa cells (Suarez et al., 2010). However, to our knowledge and until the present work, no CIS effectors (T6SS or eCIS) targeting eukaryotic cells are yet described that possess nuclease activity.

One group of evolutionarily related CIS have been shown to mediate interactions with diverse eukaryotic organisms including amoeba, grass grubs, wax moths and wasps (Böck et al., 2017; Hurst et al., 2007; Penz et al., 2012, 2010; Sarris et al., 2014; Yang et al., 2006). We recently described a related eCIS mediating the beneficial relationship between the Gram-negative bacterium *Pseudoalteramonas luteoviolacea* and a marine tubeworm, *Hydroides elegans*, hereafter *Hydroides* (Shikuma et al., 2014, 2016). We called this eCIS from *P. luteoviolacea* MACs for <u>M</u>etamorphosis <u>A</u>ssociated <u>C</u>ontractile <u>s</u>tructures, because they stimulate the metamorphosis of *Hydroides* (Shikuma et al., 2014). MACs are the first CIS discovered to form arrays of phage tail-like structures composed of about 100 tails and often measure about 1 μm in diameter. While MACs provide another example of CIS-eukaryote interactions, the range of hosts targeted by eCIS like MACs as well as the identity and mode of action of effectors that mediate these interactions remain poorly understood.

To study the interaction between MACs and eukaryotic cells, we establish an *in vitro* CIS-cell line interaction model with insect and mammalian cell types. Using these systems, we identify a new MAC effector with nuclease activity that is responsible for cytotoxicity in both cell types. Our results indicate that MACs can interact with a range of host cells and a specific effector mediates killing of eukaryotic cells.

## RESULTS

### MACs kill insect cell lines *in vitro*

To study MACs from *P. luteoviolacea* and test their effect on eukaryotic cells, we focused on an insect cell line from the Fall Armyworm *Spodoptera frugiperda* (Sf9), the closest relative to *Hydroides* where established cell lines are commercially available. Upon co-incubation of purified MACs, we observed the lysis of Sf9 cells within 48 hours (Figure 1A). As a control, we included cell-free purifications from a strain lacking the MAC baseplate gene *macB*, which is unable to produce intact phage tail-like structures and multi-tailed arrays (Shikuma et al., 2014). When Sf9 cells were co-incubated with purifications from a Δ*macB* strain or extraction buffer alone, the cells remained viable (Figure 1B and C). We quantified the activity of MACs against Sf9 cells by staining dead cells using a colorimetric stain, trypan blue, or by a fluorescent dual stain, fluorescein diacetate (FDA) and propidium iodide (PI), which stains live and dead cells, respectively. Using both methods, we observed cell death when cells were exposed to wild-type MACs, while death was not observed with purifications from a Δ*macB* strain or the extraction buffer (Figure 1D-H). Filtering extract from wild-type cells through a 0.45-μm filter abolished the cell killing effect (Figure 1I), consistent with the observation that MACs form >0.45μm arrays (Shikuma et al., 2014). Our results suggest that MACs are capable of targeting and killing insect cells.

**Figure 1.**
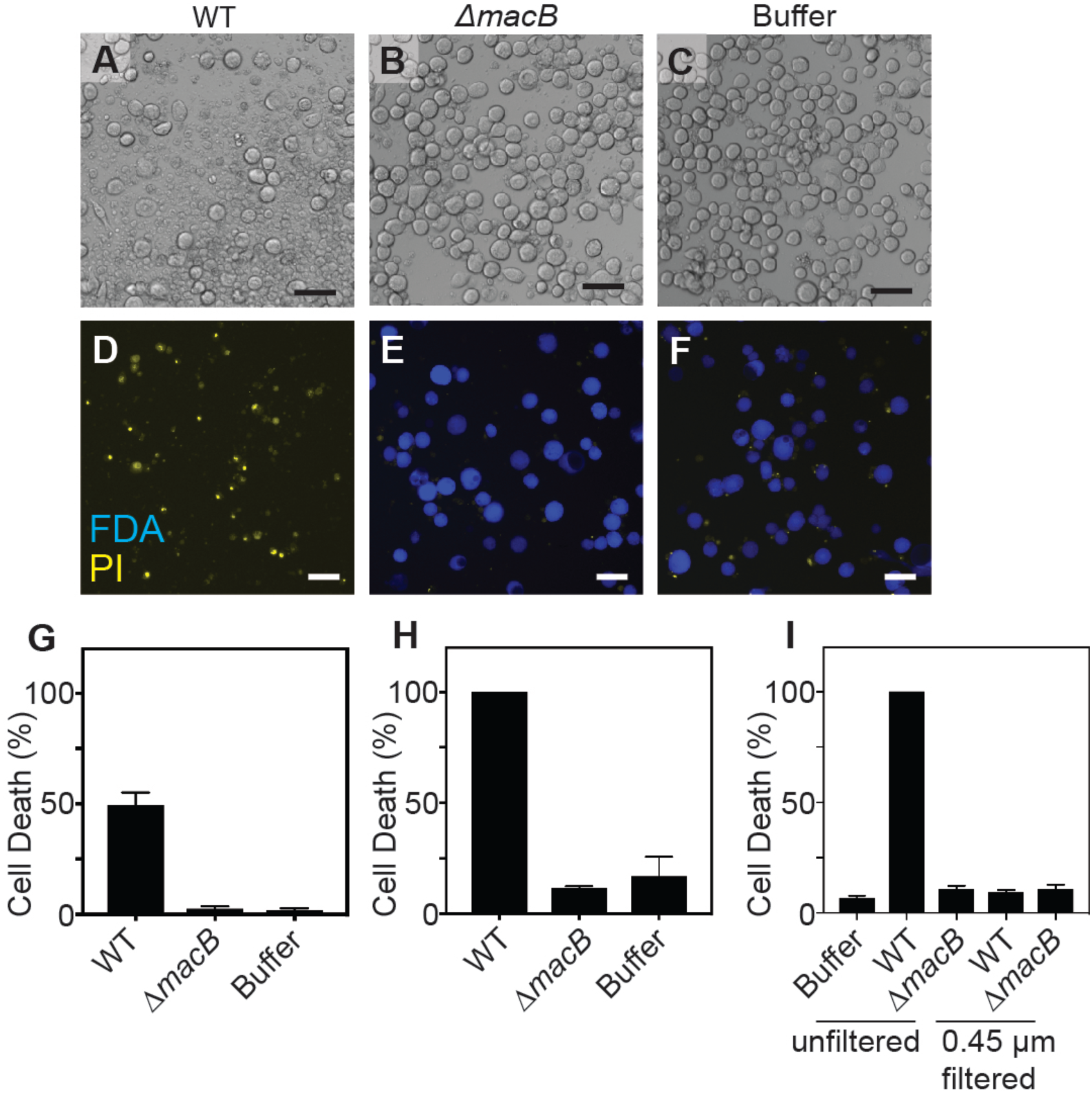
MACs cause cytotoxicity in Sf9 insect cells. (**A-C)** Sf9 cells after 48 hours incubation with MACs from wild-type *P. luteoviolacea* (WT), Δ*macB* mutant strain, or extraction buffer. **(D-F)** Live/dead staining with fluorescein diacetate (FDA) (live cells) and propidium iodide (PI) (dead cells). Scale bar is 50 μm. Quantification of cell death (%) utilizing (**G**) trypan blue and (**H**) FDA/PI live-dead stain. (**I**) Cell death of Sf9 cells exposed to MAC extract unfiltered and filtered through a 0.45 μm filter.

### Identification of a MAC effector required for killing of insect cells

We previously showed that a locus containing six genes (JF50_12590-JF50_12615) within the *P. luteoviolacea* genome is required for MACs to stimulate the metamorphosis of the tubeworm *Hydroides*, yet a mutant lacking all 6 genes is still able to produce intact MAC structures (Shikuma et al., 2016). To determine whether MACs require this same locus for killing of insect cell lines, we tested whether *P. luteoviolacea* mutants lacking each of the six genes were deficient in MAC-mediated insect cell killing. Among those six genes, only a ΔJF50_12610 mutant was unable to cause cell death upon co-incubation with insect cells (Figure 2A-G). These results were quantified and confirmed using trypan blue or FDA/PI staining (Figure 2K and L). When JF50_12610 was introduced back into its native chromosomal locus, the killing effect of MACs was restored (Figure 2H). Intriguingly, when we tested MACs from the ΔJF50_12610 mutant against larvae from *Hydroides elegans* we found that this strain was capable of stimulating metamorphosis at levels comparable to that of wild-type MACs (Figure 2M), suggesting functional structures are still present.

**Figure 2.**
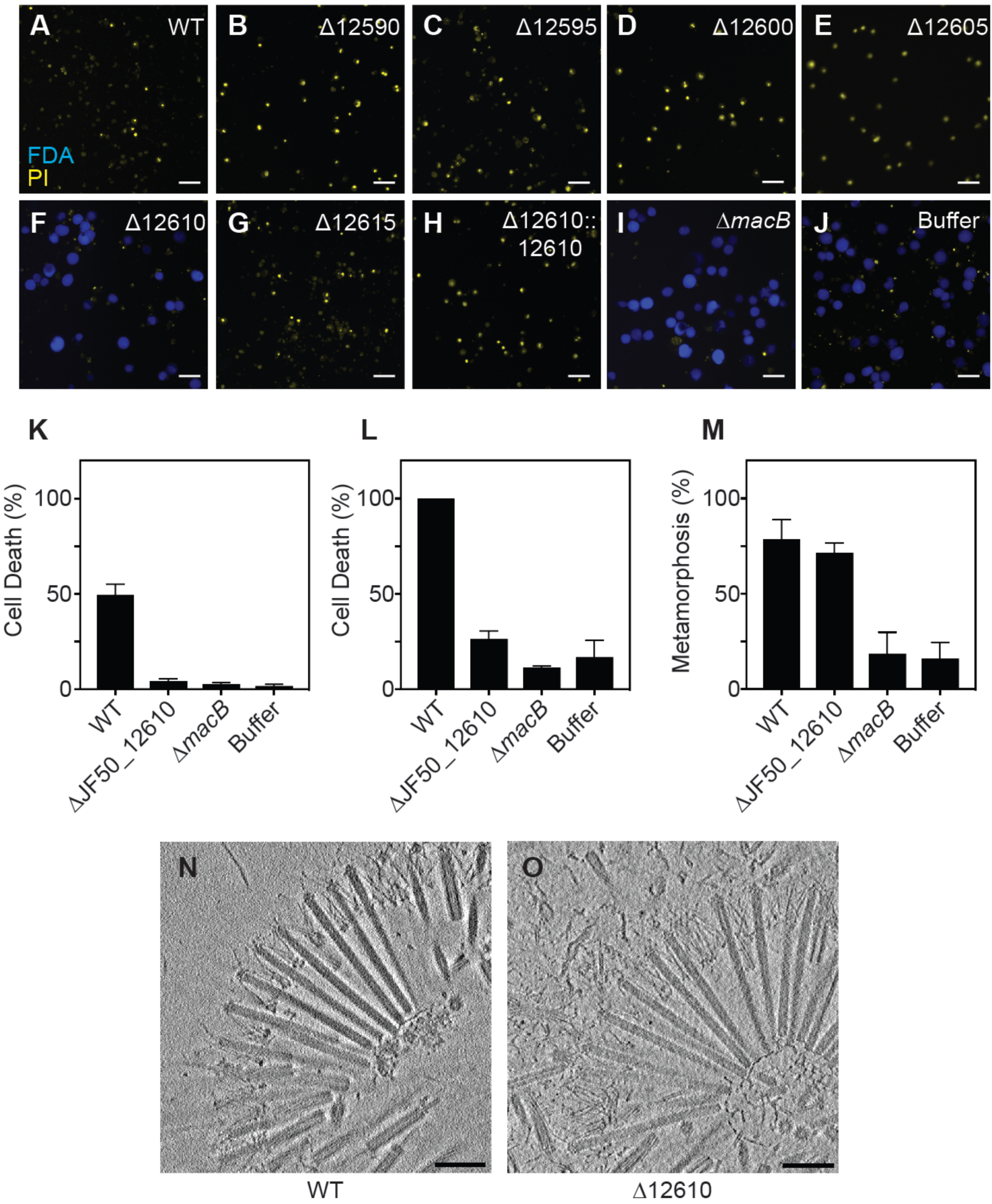
MACs require JF50_12610 to kill insect cells. (**A-J**) Sf9 cells after 48h incubation with MACs from WT, ΔJF50_12590, ΔJF50_12595, ΔJF50_12600, ΔJF50_12605, ΔJF50_12610 (Δ*pne1*), ΔJF50_12615, ΔJF50_12610∷JF50_12610, Δ*macB* strains, and extraction buffer. Scale bar is 50 μm. (**K and L**) Quantification of cell death (%) by trypan blue and FDA/PI live-dead staining. (**M)** Metamorphosis (%) of *Hydroides* larvae in response to MACs from WT, Δ*macB* or ΔJF50_12610 strains. Electron cryo-tomography images of MACs from (**N**) wild type and (**O**) ΔJF50_12610 strains showing an ordered structure with extended and contracted tubes connected by a meshwork of tail fibers. No structural differences between WT and ΔJF50_12610 arrays were observed. Shown are projections of 1.1 nm thick slices of cryotomograms. Scale bars are 100 nm.

To determine whether the ΔJF50_12610 strain produces intact MAC structures, we employed electron cryo-tomography (ECT). Upon inspection, MACs from wild type and ΔJF50_12610 were indistinguishable; forming intact phage tail-like structures in both extended and contracted conformations (Figure 2N and O). In order to confirm that JF50_12610 is part of the MAC complex, we utilized protein identification by mass spectrometry on purified MAC extracts from WT *P. luteoviolacea.* In two independent experiments, we detected JF50_12610 which indicated that the protein is associated with the MAC complex (Figure S1A). To determine whether the ΔJF50_12610 phenotype was due to the differential production of MACs, we quantified MACs tagged with super folder GFP by fluorescent microscopy and found no difference in quantity between wild-type and ΔJF50_12610 strains grown under identical conditions (Figure S1B). Our results show that JF50_12610 is required for MACs to kill insect cell lines yet does not affect the ability of MACs to stimulate tubeworm metamorphosis or the production of functional MACs. The genes within the JF50_12590-JF50_12615 locus required for tubeworm metamorphosis, and not insect cell killing, are the subject of a separate work.

### The JF50_12610 protein possesses nuclease activity *in vitro* and this activity is necessary for insect cell death

To determine the function of JF50_12610, we searched the 496 amino acid long protein for conserved domains and homologous proteins. We found that JF50_12610 contains a DNA/RNA non-specific nuclease domain (Pfam: PF13930). Analysis with the Phyre2 protein prediction program (Kelley et al., 2015) showed that residues 258-348 of JF50_12610 bear 20% identity to the nuclease Spd1 from *Streptococcus pyogenes* (Korczynska et al., 2012), and residues 267-348 bear 30% identity to the nuclease Sda1 also from *S. pyogenes* (Moon et al., 2016) (Figure 3A). Through these partial alignments, we identified a conserved glutamic acid at residue 328 corresponding to Glu164 of Spd1 and Glu225 of Sda1 that coordinate water molecules hydrating the magnesium in the enzyme’s active site (Korczynska et al., 2012; Moon et al., 2016). Consistent with its predicted function as a toxic effector against eukaryotic cells, JF50_12610 protein is predicted to contain a nuclear localization signal [NLStradamus program (Nguyen Ba et al., 2009)], which typically targets proteins to the nucleus of eukaryotic cells. Based on the predicted function of JF50_12610 and the results below, we named this effector Pne1 for *<u>P</u>seudoalteromonas* <u>n</u>uclease <u>e</u>ffector <u>1</u>.

**Figure 3.**
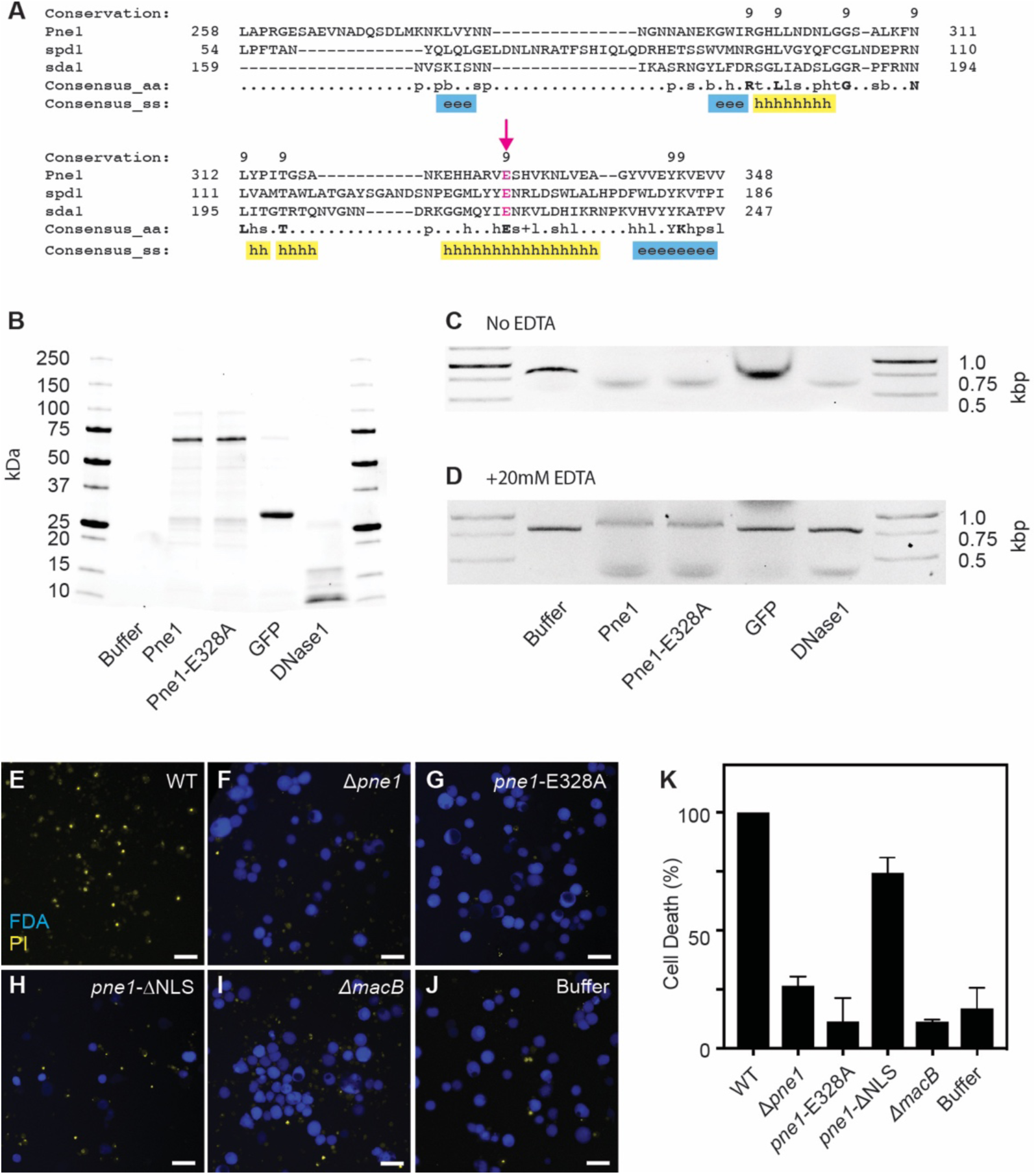
JF50_12610 (Pne1) contains a functional nuclease domain that is required for insect cell killing. (**A**) Protein alignment of Pne1 (JF50_12610), Spd1 and Sda1. Numbers indicate amino acid residues of each protein. Conserved amino acid residues indicated in bold. Consensus secondary structure (ss) alpha-helix (h, yellow) and beta-strand (e, blue). A conserved glutamic acid 328 is indicated by an arrow and highlighted in magenta. (**B**) SDS Page gel of purified wild type Pne1, Pne1-Glu328Ala, GFP, and DNase1. Representative 1% agarose gel of DNA co-incubated with Pne1, Pne1-Glu328Ala and GFP at 37°C for 2 hour (PCR product size, 0.8 Kbp) in the absence **(C)** or presence **(D)** of 20mM EDTA. Live/Dead images of Sf9 insect cells after 48 hours incubation with MACs from (**E**) WT, (**F**) Δ*pne1*, (**G**) *pne1*-Glu328A, (**H**) *pne1*-ΔNLS, (**I**) *ΔmacB*, and (**J**) MAC extraction buffer. Scale bar is 50 μm. (**K**) Quantification of cell death (%) by FDA/PI live-dead stain.

To determine whether Pne1 possesses nuclease activity, we cloned the wild type *pne1* gene and a *pne1*-Glu328Ala mutant into an IPTG-inducible vector system with N-terminal 6xHis tag and purified both proteins by nickel affinity chromatography (Figure 3B). When co-incubated with linear double-stranded DNA, Pne1 and Pne1-Glu328Ala both exhibited nuclease activity (Figure 3C), whereas a control protein, green fluorescence protein (GFP), cloned and purified under the same conditions, did not exhibit DNase activity. Based on similarities between Pne1 and magnesium-dependent homologous proteins Spd1 and Sda1, we tested the DNase activity of Pne1 protein in the presence of the divalent cation chelator, EDTA. Interestingly, Pne1 and the Pne1-Glu328Ala proteins were still functional in the presence of EDTA (Figure 3D).

To test whether Pne1 requires its nuclear localization signal or the conserved Glu328 for killing insect cells, we created *P. luteoviolacea* mutants lacking the predicted nuclear localization signal (residues 19-52) *pne1*-ΔNLS or with the *pne1*-Glu328Ala point mutation in their native chromosomal loci. Upon exposure to insect cells, MACs from the *pne1*-ΔNLS strain partially abolished the killing effect and MACs from the *pne1*-Glu328Ala strain were unable to kill insect cells when compared to wild-type MACs (Figure 3G, H, K, F). Our results show that Pne1 possesses nuclease activity *in vitro* and its nuclear localization signal may be partially responsible for the killing effect. While the Glu328Ala is not required for nuclease activity *in vitro*, this residue is necessary for MACs to kill insect cells. We are currently investigating how the Glu328Ala residue contributes to insect cell killing.

### MACs possess a broad host range, killing J774A.1 murine macrophage cell line *in vitro*

To determine whether MACs are capable of targeting a broader range of eukaryotic cells, we tested the ability of MACs to kill mammalian cells. We chose the commonly used murine macrophage cell line J774A.1 as these immune cells often encounter microbial pathogens and their effectors. Upon exposure of J774A.1 cells to wild-type MACs, we observed cell death within 24 hours (Figure 4A). In contrast, MACs from a Δ*pne1* and Δ*macB*, or buffer alone did not exhibit cell killing (Figure B-D). Quantification of cytotoxicity by lactate dehydrogenase (LDH) release assays confirmed the observed killing upon exposure to wild-type MACs and lack of killing upon exposure to MACs from the Δ*pne1* (Figure 4E). Taken together, our results show that MACs are capable of targeting and killing mammalian cells and the cell killing phenotype is dependent on the presence of Pne1.

**Figure 4.**
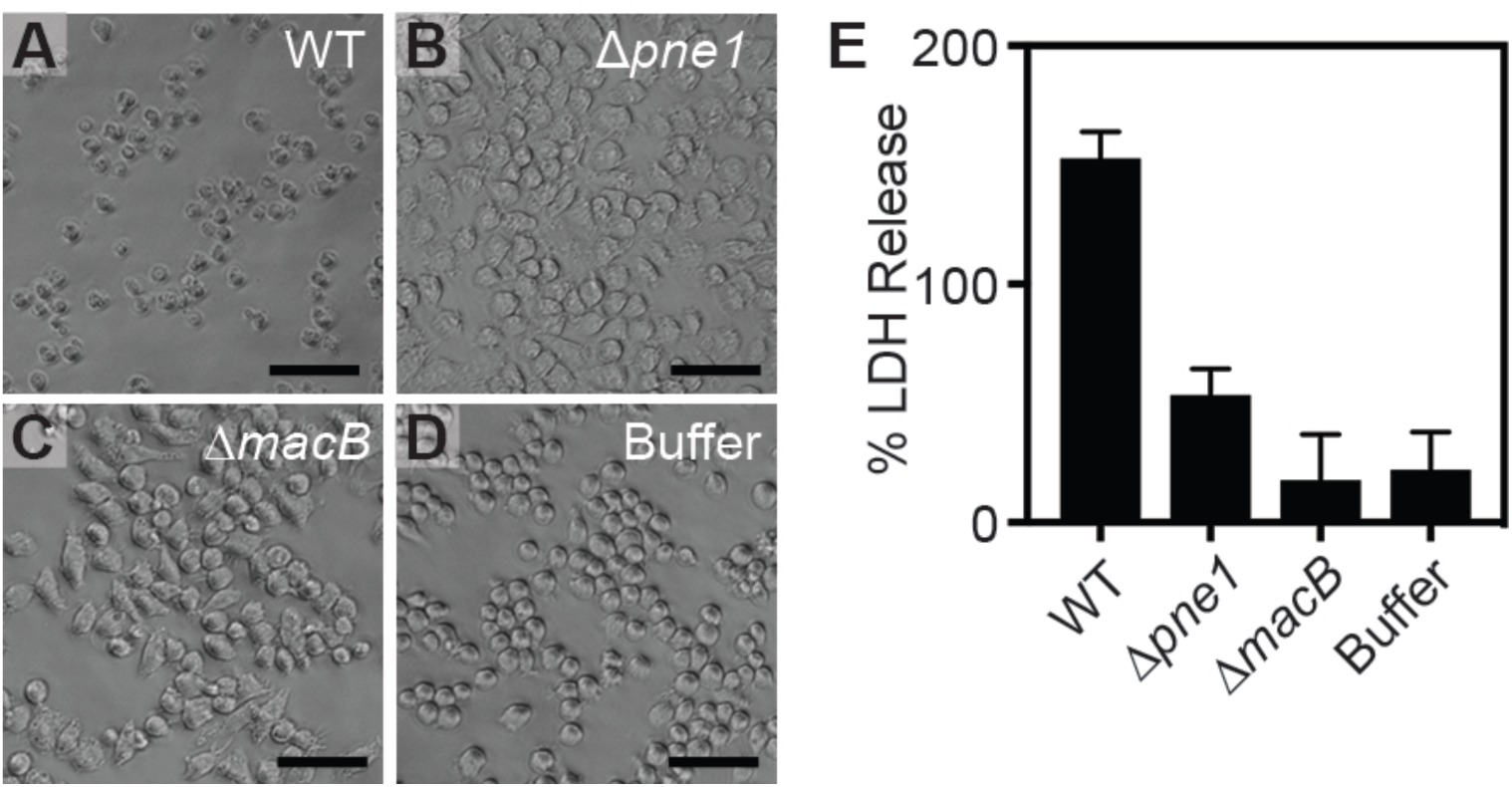
MACs kill J774A.1 murine macrophages and killing is dependent on JF50_12610. (**A-D**) J774A.1 cells after incubation with MACs from WT, ΔJF50_12610, Δ*macB* or extraction buffer. Scale bar is 50 μm. (**E**) Cell death was quantified by Lactate Dehydrogenase (LDH) release assay at 24 hours.

## DISCUSSION

In this work, we establish an *in vitro* interaction model between an eCIS and two eukaryotic cell lines. With this system, we determined that the bacterial protein, Pne1, is a novel eCIS effector and possesses nuclease activity. To our knowledge, our work is the first to identify a CIS with broad eukaryotic host range and identify the first eukaryotic-targeting CIS effector with nuclease activity.

Here, we show Pne1-dependent cell death of insect and mammalian cell lines, yet we have also previously observed that MACs stimulate metamorphosis in the tubeworm *Hydroides* (Shikuma et al., 2014). While it is unclear why MACs possess an effector-mediated killing of eukaryotic cells, some symbiotic bacteria have been shown to use CIS to modulate their host range. For example, the nitrogen-fixing plant symbiont *Rhizobium leguminosarum* limits its host range to plants in the clover family by secreting proteins through a T6SS, while mutation of the *imp* gene cluster (encoding components of the T6SS) allowed the bacterium to form functional root nodules on pea plants, normally outside of its host range (Bladergroen et al., 2003). In the environment, *Pseudoalteromonas* species are found in association with many marine invertebrates (Holmstrom and Kjelleberg, 1999), and might utilize MACs and Pne1 to antagonize specific eukaryotes, like bacterivorous ciliates, while also stimulating the metamorphosis of other eukaryotes like *Hydroides*. While several CIS nuclease effectors have been identified targeting bacterial cells, including Tde1 from the soil bacterium *Agrobacterium tumefaciens* (Ma et al., 2014), RhsA from *D. dandantii* (Koskiniemi et al., 2013), and RhsP from *Vibrio parahaemolyticus* (Jiang et al., 2018), Pne1 is the first CIS nuclease effector to our knowledge that targets eukaryotic organisms. Interestingly, the eukaryotic NLS at the N-terminus of Pne1 had a partial effect on its ability to kill insect cells, further implying its evolved role in targeting eukaryotic hosts. We are currently examining how MACs have different effects on other cell types and organisms.

The host range of eukaryotic-targeting eCISs are currently poorly understood. Intriguingly, work on related CIS show that many of them target eukaryotic organisms from diverse lineages (e.g. Grass Grubs, Wax moth, Wasps, and Amoeba)(Böck et al., 2017; Hurst et al., 2007; Penz et al., 2010; Yang et al., 2006),. While the ability of related eCIS, the Anti-feeding prophage (Afp) and *Photorhabdus* Virulence Cassettes (PVC), to target eukaryotic cells has been attributed to tail fibers that resemble receptor-binding proteins from adenovirus (Hurst et al., 2007; Yang et al., 2006), we detected no protein homology in MAC structures. We report here that MACs are capable of targeting multiple *in vitro* cell lines from insect and mammalian lineages. By further studying how eCIS, like MACs, have evolved from bacteriophage origins to target eukaryotic cells, we can begin to determine the underlying mechanisms associated with these diverse interactions.

In addition to expanding our basic understanding of CIS, our work opens the door for potentially using eCIS for biotechnology purposes. eCIS that target bacterial pathogens are already under development as narrow host-range antimicrobial agents (Scholl, 2017), for example against the gastrointestinal pathogen, *Clostridium difficile* (Gebhart et al., 2015). As syringe-like structures that deliver proteinaceous cargo to eukaryotic cells, we are currently working to develop MACs as potential delivery systems for biotechnology applications. While the mechanism of any eCIS-eukaryotic cell attachment has yet to be determined, it is tantalizing to imagine using genetically-modified CIS to deliver peptides of interest to specific eukaryotic cell types. The *in vitro* system described in this work will significantly facilitate the realization of these efforts.

## MATERIALS AND METHODS

### Bacterial Strains and Plasmid Constructions

All the strains and primers used throughout the experiments reported can be seen in tables below (Table 1, 2). All deletion and fusion strains were created according to previously published protocols (Shikuma et al., 2014, 2016). Plasmid insert sequences were verified by DNA sequencing. Deletion and insert strains were confirmed by PCR. All *E.coli* strains were grown in Luria-Bertani (LB) media at 37°C shaking at 200 revolutions per minute (RPM). All *P. luteoviolacea* cultures were grown in seawater tryptone (SWT) media (35.9 g/L Instant Ocean, 2.5g/L tryptone, 1.5 g/L yeast extract, 1.5 mL/L glycerol) at 25°C shaking at 200 RPM. Media that containing antibiotics were at a concentration of 100 mg/mL unless otherwise stated.

### MAC Purifications

MACs are produced by the marine bacteria *Pseudoalteromonas luteoviolacea* as described previously (Shikuma et al., 2014). Briefly, cells were struck out from frozen stock on SWT agar and grown at 25°C or room temperature for 1-2 days. Cells were inoculated into 5 mL SWT broth and grown for 24 hours, 25°C at 200 rpm. Cultures were inoculated 0.5 mL into 50 mL SWT in a 250 mL flask and grown for 16 hours, 25°C at 200 rpm. Cultures were transferred to a 50 mL conical centrifuge tubes and centrifuged at 4000 g for 20 minutes at 4°C. Cultures were kept on ice after this step. Subsequently, the supernatant was discarded and the pellet was resuspended in 5mL of cold extraction buffer (20mM Tris Base, 1M NaCl, 1L of deionized water, adjust the pH to 7.5 with HCl). The resuspension mixture was then transferred to a 15mL centrifuge tube and placed in the centrifuge for another 20 minutes at 4000g at 4°C. This time, the supernatant was transferred to another 15mL tube and spun down one last time with the same above conditions. After this final spin, the supernatant (now called MAC extract) was carefully poured to another clean 15mL tube and kept at 4°C until it was ready to be used for cell infections. In some experiments, MAC extract was filtered through a 0.45μm syringe filter.

### MAC Quantification

The quantification of MACs followed the same protocols and methods from with the following modifications (Thurber et al., 2009). As MACs do not contain nucleic acids, no SYBR staining was performed. We used a 0.22μm anodisc to filter clusters using the same apparatus set forth by Rower and group. The anodisc was placed on the vacuum stage with the shiny side facing up. Then, the glass column and clamp were secured on top of it and 2mL of 1XPBS was added and allowed to run through the column. Once the PBS was filtered and the anodisc apparatus was successfully placed, 100μL of MAC extract was added to 900μL of 1XPBS and run through the filter. Once all the liquid passed through the anodisc, the clamp and column were taken out before shutting off the vacuum to assure that the anodisc stayed on the stage. The anodisc was placed on a kimwipe to allow it to dry properly for a few minutes. Once dried, a slide was made using 10μL of the microscope mount (10% ascorbic acid 1XPBS with 50% glycerol, filter sterilize mount with 0.02μm filter) “sandwiching” the anodisc. A ZEISS microscope was used to quantify MACs tagged with sf-GFP. Quantification of MACs were performed using column diameter, field of view and volume.

### Insect Cell Culture

To test the effect of *P. luteoviolacea* MACs interaction with eukaryotic insect cell lines, we used Sf9 cells (Novagen #71104-3). Cells were cultured in ESF 921 Insect Cell Culture Medium (Expression systems #96-001-01). Frozen cell line stocks were taken from −80°C freezer and thawed quickly at 27°C in a water bath. Thawed cells (~1mL) were added to a 125mL flask containing 10mL of room temperature ESF 921. Cells were then placed in a 27°C incubator shaker, shaking at 130 rpm, in the dark. Cell lines were maintained by passaging every 2-3 days using routine cell culture techniques.

### Insect Cell Co-incubation with MACs

Prior to treatment with MACs, Sf9 cells were passaged five to twenty times and seeded into 24-well tissue culture plates at a density of 4×10^5^ cells/mL a few hours before infection. MACs that were extracted no more than a week prior to cell infections were added to each well of the 24-well plates with cells at a 1:50 ratio (10μL of the extracts into 500μL of cells). Infected cells were incubated at 27°C, adherently. At 48 hours, trypan blue or PI/FDA was added to each well of the 24-well infection plates and microscopic images were taken of each well to visualize and quantify cell viability.

### Protein Purification

The JF50_12610 wild type, JF50_12610-Glu328Ala and Green Fluorescence Protein (GFP) were cloned into a pET15b vector with N-terminal 6xHis tag. The protein was overexpressed in 500mL of LB medium growing at 37°C until 0.9OD_600_ and induced using 1.0mM IPTG at 25°C for 4 hours, followed by a centrifugation step of 4000g for 20min at 4°C. The cell pellet was resuspended using a native lysis buffer (0.5M NaCl, 20mM Tris-HCl, 20mM imidazole, pH 8). The resuspended cell lysate was then submitted to a French press 2X, which was followed by sonication on high (10 seconds sonication, repeated twice, done on ice). Following cell sonication, the supernatant was purified by a bulk Ni-NTA beads, washed with 20mL lysis buffer on column, and eluted using elution buffer (0.5M NaCl, 20mM Tris-HCl, 250mM imidazole, pH 8) where protein fractions were collected. Eluted protein was buffer exchanged using pierce protein concentrators (thermos fisher cat# 88513) into (0.15M NaCl, 20mM Tris-HCl, pH 8) Collected protein fractions were then quantified using a Thermo-Fisher Pre-diluted protein standards kit (catalog# 23208), and subsequently read on a Biotek plate reader (catalog# 49984). Proteins were then normalized to equal concentrations prior to nuclease assay.

### Nuclease Assay

In order to test the bioinformatically predicted nuclease activity found in JF50_12610, a DNase assay was developed. The wild type JF50_12610, JF50_12610-E328A and Green Fluorescence Protein (GFP) were purified simultaneously and identically prior to assay. After protein purification, protein concentrations were normalized to 0.5μg/μL, including a positive control commercially available DNase I (BioBasic #DD0099-1). All normalized proteins were then incubated for 2 hours at 37°C with a PCR DNA fragment of known size and concentration and NEB cutsmart buffer (New England Biosciences #B72045). A reaction total volume of 20μL consisted of 12μL of protein, 6μL of DNA diluted in buffer (0.15M NaCl, 20mM Tris-HCl, pH 8), and 2μL of 10X cutsmart buffer. After 1-hour of incubation, all reactions and their replicates were resolved in an 1% agarose gel stained with EtBr in 1XTBE (Tris-Borate EDTA, BioBasic #A00265) at 110 volts for about 45 minutes. Gel products were then visualized in a BioRad gel imaging machine (Gel Doc™ EZ System #1708270) appropriately. The +EDTA reaction was completed identically to the protocol above with the exclusion of the NEB cut smart buffer and addition of 20mM EDTA (final concentration) to the reaction mixture.

### Gentle MAC extraction and mass spectrometry

*P. luteoviolacea* was grown in 50mL Marine Broth (MB) media in 250mL flasks at 30°C for 6 hours or overnight (12-14 hours). Cells were centrifuged for 30 minutes at 7000 g and 4°C and resuspended in 5mL cold extraction buffer (20 mM Tris, pH 7.5, 1M NaCl). The resuspensions were centrifuged for 30 minutes at 4000 g and 4°C and the supernatant was isolated and centrifuged for 30 minutes at 7000 g and 4°C. The pellet was resuspended in 20-100μL cold extraction buffer and stored at 4°C for further use. All mass spectrometry was done by the Functional Genomics Center Zurich (FGCZ). To prep the MAC extracts for mass spectrometry, the extracts were precipitated by mixing 30μL of sample with 70μl water and 100μl 20% TCA. The samples were then washed twice with cold acetone. The dry pellets were dissolved in 45μl buffer (10mM Tris/2mM CaCl_2_, pH 8.2) and 5μl trypsin (100ng/μL in 10mM HCl). They were then microwaved for 30 minutes at 60°C. The samples were dried, then dissolved in 20μL 0.1% formic acid and transferred to autosampler vials for LC/MS/MS. 1μL was injected.

### Electron cryo-tomography of MAC arrays

For ECT imaging of MAC arrays, *P. luteoviolacea* WT and ΔJF50_12610 were cultured for 5-6 hours at 30°C and subsequently centrifuged for 30 minutes at ~7000 g. The pellet was resuspended in 5mL of extraction buffer and centrifuged for 30 minutes at 4000 g to separate intact cells from MAC arrays. The supernatant was carefully transferred into a new tube and centrifuged for 30 minutes at 7000 g. The pellet was resuspended in about 50 μL of extraction buffer and mixed with Protein A-conjugated 10-nm colloidal gold (Cytodiagnostics Inc.) before plunge freezing. Plunge freezing and ECT imaging were performed according to Weiss et al. (Weiss et al., 2017). 4 μL of sample was applied to glow-discharged EM grids (R2/1 copper, Quantifoil), blotted twice from the back for 3.5 s and vitrified in liquid ethane–propane using a Vitrobot Mark IV (ThermoFisher). ECT data were collected on a Titan Krios (ThermoFisher) transmission electron microscope equipped with a Quantum LS imaging filter (Gatan, slit with 20 eV) and K2 Summit direct electron detector (Gatan). Tilt series were acquired with the software SerialEM (Mastronarde, 2005) using a bidirectional tilt scheme. The angular range was −60° to +60° and the angular increment was 2°. The total electron dose was between 60–100 electrons per Å^2^ and the pixel size at specimen level was 2.72 Å. Images were recorded in focus with a Volta phaseplate (ThermoFisher) for WT and without phaseplate at 5 μm under-focus for ΔJF50_12610. Tilt series were aligned using gold fiducials and three-dimensional reconstructions were calculated by weighted back projection using IMOD (Mastronarde, 1997). Visualization was done in IMOD. Contrast enhancement of the ΔJF50_12610 tomogram was done using the tom_deconv deconvolution filter (https://github.com/dtegunov/tom_deconv).

### Macrophage Cell Culture

To test the effect of *P. luteoviolacea* MACs on eukaryotic cells, we tested their effect on murine macrophages J774A.1. Macrophage cells were cultured in DMEM (Dulbecco’s Modified Eagle’s Medium, Gibco #10566-016), which was prepared with the addition of 10% of heat-inactivated fetal bovine serum (FBS) and 1% sodium pyruvate. Frozen cell line stocks were taken from nitrogen tank and thawed quickly at 37°C in a water bath. Thawed cells (~1mL) were added to a 50mL conical centrifuge tube containing 5mL of pre-warmed DMEM. Cells were then centrifuged at 500 g at room temperature for 4 minutes. The supernatant was carefully discarded, and cells were resuspended in 7.5mL of pre-warmed DMEM. This step assured complete removal of DMSO from frozen cell stocks. Resuspended cells were transferred from the 50mL conical centrifuge tube into a 25cm^2^ tissue culture flask and incubated at 37°C with 5% CO_2_. Cell lines were maintained by passaging every 2-3 days using routine cell culture techniques.

### Macrophage Cell Co-incubation with MACs and LDH Assay

Prior to exposure to MACs, J774A.1 cells were passaged two to five times and seeded into 24-well tissue culture plates at a density of 4×10^5^ cells/mL the night before the treatment. MACs that were extracted no more than a week prior to cell treatment were added to each well of the 24-well plates with cells at a 1:50 ratio (10μL of the extracts into 500μl of cells). Treated cells were incubated at 37°C with 5% CO_2_. Culture supernatants were collected at between 1 and 24 hours post-infection. At each time point, the plate was first spun down at 400 g for 2 minutes, then 100μL of the supernatant was transferred into a 96-well plate in duplicate. For t=0 time point, remaining supernatants were removed and cells were lysed with PBS+ 1% tritonX. 100 μL of the cell lysates was also transferred to 96-well plates. Microscopic images were taken of each well to visualize and record the cells viability, using a phase-contrast microscopy.

To assess cell lysis, we quantified the release of lactate dehydrogenase (LDH) using Takara LDH Cytotoxicity Detection Kit (#MK401). The LDH reaction mix was made per manufacturer’s instructions and kept away from the light. Cell supernatants were diluted 1:10 in PBS and diluted supernatants were mixed 1:1 with the LDH reaction mix. After the addition of the reaction mix, plates were incubated for 30 minutes at room temperature away from the light and absorbance was measured at 492nm using a plate reader. Total LDH level at t=0 was calculated by adding LDH levels of t=0 supernatants and lysates. LDH release 24 hours post-treatment was calculated as a fraction of total LDH available at the time of infection (t=0).

## ACKNOWLEDGEMENTS

We thank Shari Wu and Tom Huxford for kindly providing us with insect cell lines and culturing technique guidance. We also thank Saori Amaike Campen and Nicole Yee from JCVI for their help in the maintenance of J774A.1 macrophage cell lines. We acknowledge the imaging facility ScopeM for instrument access at ETH Zürich. This work was supported by the funds provided by Office of Naval Research (N00014-17-1-2677 to N.J.S. and S.B.), Office of Naval Research (N00014-16-1-2135 to N.J.S), Alfred P. Sloan Foundation, Sloan Research Fellowship (to N.J.S), San Diego State University (to N.J.S), J. Craig Venter Institute (to S.B.), European Research Council (to M.P.), Swiss National Science Foundation (to M.P.) and Gebert Rüf Foundation (to M.P.).

