## Supplemental Information for "Bacterial Phage Tail-like Structure Kills Eukaryotic Cells by Injecting a Nuclease Effector"

**Table 1. Table of Strains.** List of strains produced and used in this work.

| Strain No. | Strain | Genotype | Source |
|---|---|---|---|
| NJS 002 | HI1 Str$^R$ | *P. luteoviolacea* HI1, Str$^R$ | (Huang et al., 2012) |
| NJS 213 | Δ*macB* | *P. luteoviolacea* HI1 Δ*macB*, Str$^R$ | (Shikuma et al., 2014) |
| NJS 235 | ΔJF50_12590-F50_12615 | *P. luteoviolacea* HI1, Str$^R$ ΔR4 | (Shikuma et al., 2016) |
| NJS 289 | ΔJF50_12590 | *P. luteoviolacea* HI1, Str$^R$ ΔJF50_12590 | This Study |
| NJS 287 | ΔJF50_12595 | *P. luteoviolacea* HI1, Str$^R$ ΔJF50_12595 | This Study |
| NJS 285 | ΔJF50_12600 | *P. luteoviolacea* HI1, Str$^R$ ΔJF50_12600 | This Study |
| NJS 283 | ΔJF50_12605 | *P. luteoviolacea* HI1, Str$^R$ ΔJF50_12605 | This Study |
| NJS 281 | ΔJF50_12610 | *P. luteoviolacea* HI1, Str$^R$ ΔJF50_12610 | This Study |
| NJS 279 | ΔJF50_12615 | *P. luteoviolacea* HI1, Str$^R$ ΔJF50_12615 | This Study |
| NJS 373 | ΔJF50_12600::12610 | *P. luteoviolacea* HI1, Str$^R$ ΔJF50_12610::12610 | This Study |
| NJS 444 | JF50_12610-E328A | *P. luteoviolacea* HI1, Str$^R$ JF50_12610-E328A | This Study |
| NJS 209 | *macB*-GFP | *P. luteoviolacea* HI1 *macB-GFP*, Str$^R$ | (Shikuma et al., 2014) |
| NJS 380 | JF50_12610-ΔNLS | *P. luteoviolacea* HI1 12610-ΔNLS, Str$^R$ | This Study |
| **Plasmid No.** | **Plasmid** | **Genotype** | **Source** |
| pNJS 007 | pCVD443 | Amp$^R$, Km$^R$, sacB, pGP704 derivative | (Huang et al., 2012) |
| pNJS 266 | pCVD443_Δ12590 | pCVD443::Δ12590 Amp$^R$ , Km$^R$ | This Study |
| pNJS 265 | pCVD443_Δ12595 | pCVD443::Δ12595 Amp$^R$ , Km$^R$ | This Study |
| pNJS 264 | pCVD443_Δ12600 | pCVD443::Δ12600 Amp$^R$ , Km$^R$ | This Study |
| pNJS 263 | pCVD443_Δ12605 | pCVD443::Δ12605 Amp$^R$ , Km$^R$ | This Study |
| pNJS 262 | pCVD443_Δ12610 | pCVD443::Δ12610 Amp$^R$ , Km$^R$ | This Study |
| pNJS 261 | pCVD443_Δ12615 | pCVD443::Δ12615 Amp$^R$ , Km$^R$ | This Study |
| pNJS 433 | pCVD443_Δ12610::12610 | pCVD443::12610 Amp$^R$ , Km$^R$ | This Study |
| pNJS 545 | pCVD443_12610-E328A | pCVD443_12610-E328A Amp$^R$ , Km$^R$ | This Study |
| pNJS 435 | pCVD443_12610-ΔNLS | pCVD443_12610-ΔNLS Amp$^R$ , Km$^R$ | This Study |
| pNJS 430 | pET15b_12610 | pET15b_12610 Amp$^R$ | This Study |
| pNJS 544 | pET15b_12610-E328A | pET15b_12610-E328A Amp$^R$ | This Study |
| pNJS 398 | pET15b_GFP | pET15b_GFP Amp$^R$ | This Study |

**Table S2. Primers used in this work.**

| Primer | Sequence |
|---|---|
| *macB*_dA | GGTCGACGGATCCCAAGCTTCTTCTAGAGGTACCGCATGCAACCCAGACACTGAGGTGCT |
| *macB*_dB | TTTCCATTTTCCAATCCCTTCGCCAGAGATAAGTGATTGACTACGA |
| *macB*_dC | TCGTAGTCAATCACTTATCTCTGGCGAAGGGATTGGAAAATGGAAA |
| *macB*_dD | ACACAACGTGAATTCAAAGGGAGAGCTCGATATCGCATGCCATAACCTGGCTGAGCACCT |
| 1556_dA | TGATGGGTTAAAAAGGATCGATCCTCTAGATTGGAGCAATAAACGGGTTC |
| 1556_dB | GTTCATAATTAAACTGCGATCGCAGCCATAAGGCCTCCTTGATA |
| 1556_dC | TATCAAGGAGGCCTTATGGCTGCGATCGCAGTTTAATTATGAAC |
| 1556_dD | TTTTGAGACACAACGTGAATTCAAAGGGAGAGCTCCGCTTTGGGTACTGGCTTTA |
| 1556_intF | CCGAGCAAACGTTATCACAA |
| 1556_intR | TCAGCGCTCTCATTATGTGC |
| 1555_dA | TGATGGGTTAAAAAGGATCGATCCTCTAGACCGAGCAAACGTTATCACAA |
| 1555_dB | CCTTGCATGAGGTTAAGAAAGTTTGACGTACCCTTCAGCCATATT |
| 1555_dC | AATATGGCTGAAGGGTACGTCAAACTTTCTTAACCTCATGCAAGG |
| 1555_dD | TTTTGAGACACAACGTGAATTCAAAGGGAGAGCTCGATGCGGTAACGGTTGTTCT |
| 1555_intF | AGCGATTGATGCTGAACAAA |
| 1555_intR | ACCATCGCATAACCCGTAAC |
| 1554_dA | TGATGGGTTAAAAAGGATCGATCCTCTAGATACGCCGTCCAGTTAGGACT |
| 1554_dB | GTTTGTTAACGTCACGGCAGCTGCATTGCCATTTAAACTCC |
| 1554_dC | GGAGTTTAAATGGCAATGCAGCTGCCGTGACGTTAACAAAC |
| 1554_dD | TTTTGAGACACAACGTGAATTCAAAGGGAGAGCTCATTGATTGGAAGCGCGATAG |
| 1554_intF | TTTATGAGGCACCAACGACA |
| 1554_intR | GCCTGTGCCGTTTTATCTGT |
| 1553_dA | TGATGGGTTAAAAAGGATCGATCCTCTAGAGGCGATCAGTGGAGTGAAGT |
| 1553_dB | AATACTTCTTGCTCAGCCCCGCGTGCTTCTTCTGTCATGT |
| 1553_dC | ACATGACAGAAGAAGCACGCGGGGCTGAGCAAGAAGTATT |
| 1553_dD | TTTTGAGACACAACGTGAATTCAAAGGGAGAGCTCTCAGAACCAGCAGTCTCACG |
| 1553_intF | CGGGCCTAGAAATCACTCAA |
| 1553_intR | TCGACGTCAAATCAGTCGAG |
| 1552_dA | TGATGGGTTAAAAAGGATCGATCCTCTAGAGAGAGCAAGAAGTGGCGAGT |
| 1552_dB | TAGCCTTTTAGTGCCGCTTTTGAGGCGTCCATATCTGACA |
| 1552_dC | TGTCAGATATGGACGCCTCAAAAGCGGCACTAAAAGGCTA |
| 1552_dD | TTTTGAGACACAACGTGAATTCAAAGGGAGAGCTCTGCTGACCAAGCAGATTGAC |
| 1552_intF | GGGCAATTGTTGTGGATTTT |
| 1552_intR | TGATCCCAAACCACTTGTGA |
| 1551_dA | TGATGGGTTAAAAAGGATCGATCCTCTAGAGACTGCTGGTTCTGATTCGAT |
| 1551_dB | AACAGATCATTACATTAAAATGAGCCTCTGTTCTTGTTGTTGCATTTCA |
| 1551_dC | TGAAATGCAACAACAAGAACAGAGGCTCATTTTAATGTAATGATCTGTT |
| 1551_dD | TTTTGAGACACAACGTGAATTCAAAGGGAGAGCTCCTTCTCCATTTTCGCCTTTG |
| 1551_intF | CGTTTTCAGTGACCATCACG |
| 1551_intR | CGGTGGGCAAAAAGGTATAA |
| pET15b_12610_F1 | CCTGGTGCCGCGCGGCAGCCATATGATGATGTCAGATATGGACGC |
| pET15b_12610_R1 | TCGGGCTTTGTTAGCAGCCGGATCCTTAGCCTTTTAGTGCCGCTT |
| 12610-ΔNLS_B | ATACTCATATCTGAGTTTTGTTGCTTCTGCTCTTCGCTAT |
| 12610-ΔNLS_C | ATAGCGAAGAGCAGAAGCAACAAAACTCAGATATGAGTAT |
| JF50_12610_SeqF1 | CCTGAAGGGTCGTTTTCAGT |
| JF50_12610_E328A_F | GCTCGAGTTGCGTCTCATGTA |
| JF50_12610_E328A_R | TACATGAGACGCAACTCGAGC |

**A**

| | | WT 1 | WT 2 |
|---|---|---|---|
| JF50_12610 | Spectral count<br>Coverage | 2<br>5.0% | 3<br>7.3% |
| Hcp1 | Spectral count<br>Coverage | 35<br>93% | 49<br>97% |

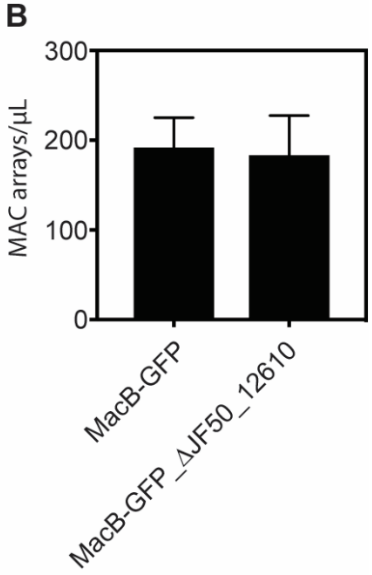


**Figure S1. JF50_12610 is associated with the MAC complex and a JF50_12610 deletion produces the same quantity of intact MACs as wild type.** (**A**) Table summarizing mass spectrometry data that indicates that JF50_12610 (Pne1) is associated with the MAC complex. Counts of the Hcp1 (MAC tail tube) protein are provided as a control. (**B**) Quantification of MAC production between wild type and ΔJF50_12610 strains. Both strains were tagged with a super folder-GFP on the baseplate as described previously (Shikuma et al., 2014). The number of MAC arrays were quantified between wild type and ΔJF50_12610 mutant using a hemocytometer and fluorescence microscopy.
